# Sweeps in time: leveraging the joint distribution of branch lengths

**DOI:** 10.1101/2021.01.27.428367

**Authors:** Gertjan Bisschop, Konrad Lohse, Derek Setter

**Affiliations:** Institute of Evolutionary Biology, University of Edinburgh, UK

## Abstract

Current methods of identifying positively selected regions of the genome are limited by their underlying model in two key ways: the model cannot account for the timing of the adaptive event and the analytic predictions are limited to single nucleotide polymorphisms. Here we develop a tractable method of describing the effect of positive selection on the genealogical histories in the surrounding genome, explicitly modeling both the timing and context of the adaptive event. In addition, our framework allows us to go beyond simple polymorphism data. We are able to leverage information contained in patterns of linked variants, and even with very small sample sizes, our analytic framework has high power to identify historically adaptive regions of the genome and to correctly infer both the time and strength of selection. Finally, we derived the marginal distribution of genealogical branch lengths at a locus affected by selection acting at a linked site. This provides a much-needed link between current theoretical models to recent advances in simulation procedures that have allowed researchers both to examine the evolution of genealogical histories at the level of full chromosomes and build methods that attempt to reconstruct full ancestries from genome sequence data.

## 1 Introduction

The variation that we observe in genome sequence data is the result of the combined demographic and selective forces acting in the evolutionary history of a population. While demography shapes genetic variation uniformly throughout the genome, natural selection has localized effects on genetic variation near the targets of past selection. Recombination attenuates the strength of this effect with increasing distances from any selected site. Despite this key difference, distinguishing the signatures of natural selection from those of demography in genomic variation remains a significant challenge.

Substantial effort has been made to describe the effect of positive selection on the genealogical history at linked neutral sites and to develop methods to detect the footprint of adaptive evolution in genomic data (for an overview, see Hejase et al. (2020a)). Here we focus on the class of parametric model-based methods that identify the signature of hard selective sweeps as a local distortion of ancestry caused by genetic hitchhiking (for a survey of such methods, see Pavlidis and Alachiotis (2017)). When a new adaptive mutation sweeps through a population, the hitchhiking of linked neutral variation leads to a local reduction in genetic diversity (Maynard Smith and Haigh, 1974) and generates a statistically detectable footprint in the site-frequency spectrum (SFS) (Kim and Stephan, 2002). This forms the basis for a number of composite likelihood methods to detect selective sweeps such as SweepFinder (Nielsen et al., 2005), SweepFinder2 (DeGiorgio et al., 2016), SweeD (Pavlidis et al., 2013), and for adaptive introgression sweeps, VolcanoFinder (Setter et al., 2020).

However, many of these methods are limited in at least three fundamental ways. Firstly, their focus on summaries of average diversity and divergence discards relevant information in the co-occurrence of closely linked variants. Secondly, assuming equilibrium population dynamics has been shown to increase both false positive and false negative error rates (Crisci et al., 2013). Finally, current sweep scanning approaches assume that the population has been sampled at the time of fixation of the beneficial mutation, leading to a decrease in power to detect increasingly old sweeps. Given these limitations, it remains an open question how much additional information about old selective sweeps is contained in sequence variation.

### 1.1 Approximating Sweeps

Since the introduction of the hitchhiking model (Maynard Smith and Haigh, 1974), many approximations for the effect of a selective sweep have been developed using the coalescent framework of (Kingman, 1982; Hudson, 1983; Tajima, 1983). Here, the fixation of a new beneficial mutation has the effect of genetically structuring the ancestry at linked neutral loci (Kaplan et al., 1989a; Stephan et al., 1992; Barton et al., 2004). During the sweep, coalescence can only occur among lineages on the same genetic background as the selected locus, while recombination may move lineages from the selected onto a neutral genetic background. However, analytic expressions to quantify these effects are only possible with further simplifications of the model. The genome scanning methods mentioned above are based on the star-like approximation for the selective sweep (Barton, 1998, 2000; Durrett and Schweinsberg, 2005; Berg and Coop, 2015), as it is relatively accurate, yet computationally tractable for large sample sizes. We can view the star-like approximation as follows: assuming selection is strong (*N_e_s* >> 1), fixation of the beneficial mutation happens nearly instantaneously on the coalescent time scale. Lineages either recombine out of the sweep individually or coalesce in a single multiple-merger event at the origin of the beneficial mutation. However, this assumption fails either when selection is weak or at intermediate recombination distances from the selective target when selection is strong, and this leads to biased parameter estimates (Barton, 1998; Santiago and Caballero, 2005; Setter et al., 2020; Hartfield and Bataillon, 2020; Charlesworth, 2020).

More accurate approximations for the effect of selective sweeps on genealogies have been developed (Bossert and Pfaffelhuber, 2013), which, although more cumbersome mathematically, may avoid biases in parameter estimates and genome scans. The initial growth of a beneficial mutation behaves like a super-critical branching process (Evans and O’Connell, 1994; Kaplan et al., 1989a; Barton, 1998). Conditioned on fixation, the stochastic increase in frequency is well-approximated by a pure-birth or Yule process. The structured co-alescent that describes the genealogy at a linked neutral locus is then well approximated by marking the lineages in the Yule tree by recombination events (Schweinsberg and Durrett, 2005; Etheridge et al., 2006; Pfaffelhuber et al., 2006). Thus, in contrast to the star-like approximation, lineages on the selected background are assumed to coalesce pairwise during the sweep and can later recombine out of the sweep. Modelling the sweep phase as a time interval during which the coalescent is governed by the Yule process (Hermisson and Pfaffelhuber, 2008) is possible for reasonably strong selection and small sample sizes. We will refer to this as the full Yule approximation. An alternative approach, which extends to larger samples, is to use the Yule process to derive better approximations for a model that assumes that the sweep is instantaneous (on the coalescent time scale) (Etheridge et al., 2006; Bossert and Pfaffelhuber, 2013). The sampling formulae derived by Bossert and Pfaffelhuber (2013) assume that a sweep partitions lineages at a linked neutral locus into three families: non-recombining lineages, early-recombining lineages and late recombinant lineages. We will refer to this as the instantaneous Yule approximation. Like the star-like approximation, the instantaneous Yule approximation is an instantaneous partitioning of the sample, but allows for up-to two multiple-merger events. (Pfaffelhuber et al., 2006).

### 1.2 Overview

The motivation of the present study is to develop a full analytic description of the effect of a hard selective sweep occurring on the distribution of genealogies at nearby neutral sites and to explore how this can be used to improve likelihood based inference.

The paper is structured as follows: First, we briefly summarize approximate models of selective sweeps and show how a hard selective sweep occurring at an arbitrary time in the past can be embedded in the generating function (GF) for the distribution of the genealogy introduced by (Lohse et al., 2011). The GF provides a recursive description of the full genealogy of a sample for a general class of structured coalescent processes with discrete events. While previous applications of the GF have focused on models of demographic history (Bunnefeld et al., 2015; Lohse et al., 2016), here we use the GF framework to describe the genealogy at a neutral locus associated with a hard sweep occurring at a given time in the past.

Secondly, we use the GF to derive (and re-derive) analytic predictions for the effect of a sweep on mean genetic diversity, the SFS, and the probability of genealogical topologies in the vicinity of a sweep target. In addition, we obtain the marginal distributions of the length of branches with *i* descendants among the sample (i-Ton branches) which underlie the SFS, and we extend the model to the adaptive introgression scenario of Setter et al. (2020).

Finally, to connect these results to sequence data, we use the GF method to compute the probability distribution of block-wise configurations of completely-linked mutations in the region of the genome affected by the selective sweep and develop a simple composite likelihood framework based on the blockwise SFS (bSFS). We use comparisons to forewards simulations (Haller and Messer, 2019) throughout to quantify the robustness of our analytic predictions, compare the accuracy of the star-like and the instantaneous Yule approximation and assess the power of our composite likelihood method to jointly estimate the sweep time and the strength of selection.

## 2 Model and Methods

### 2.1 Evolutionary History

We consider *n* lineages sampled from a panmictic population of N diploid individuals that evolves according to a Wright-Fisher model. We initially assume that each lineage is uniquely labelled, i.e. the data are polarized relative to an outgroup and each haplotype is fully phased (we relax these assumptions when considering inference). In Fig 1, we uniquely label the lineages ancestral to each sampled individual *a, b, c, d, e,* and *f*. A coalescence event may generate, e.g. branch type bc which is ancestral to lineages *b* and *c*.

**Figure 1:**
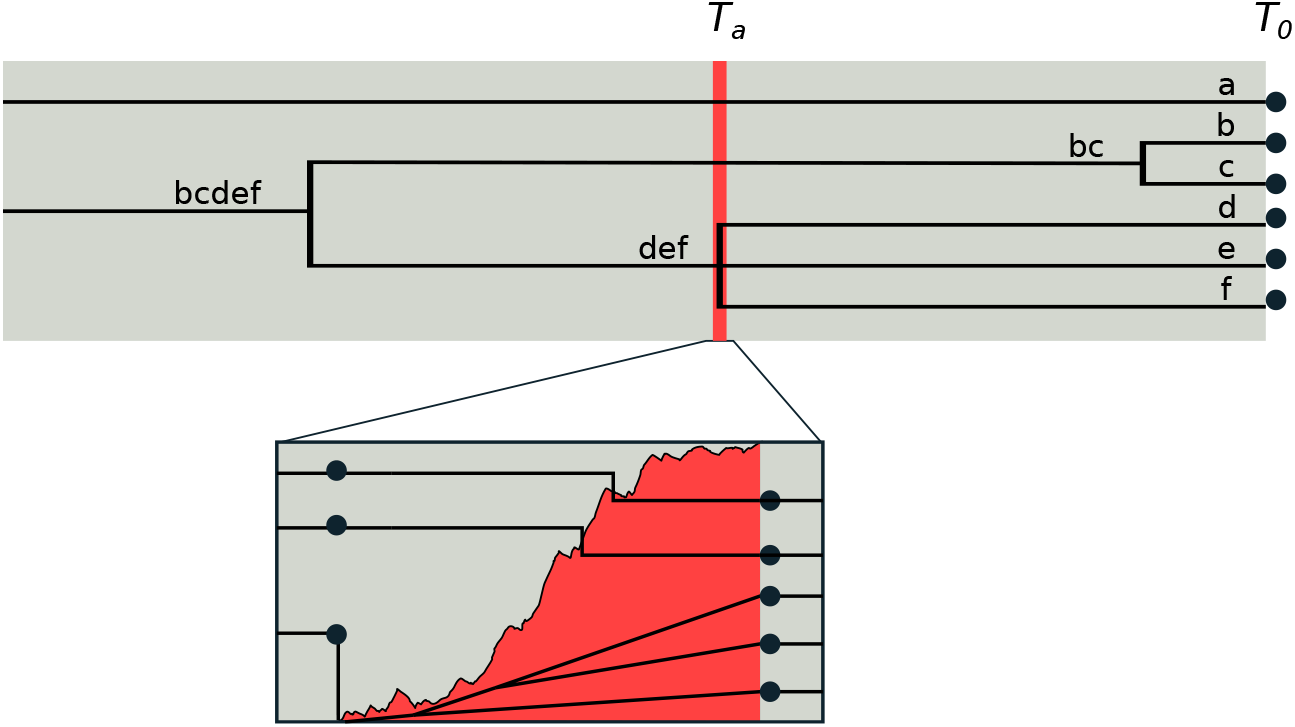
Model. The effect of an old selective sweep at time *T_a_* on a sample of six lineages {*a, b, c, d, e, f*} at a nearby neutral site. Tracing the genealogy pastward, we first observe a neutral coalescence of the *b* and *c* lineages. The second event is the selective sweep, which occurs quickly on the time scale of coalescent events. This induces what appears to be a multiple-merger coalescence of *d, e*, and *f* (as in the star-like approximation). On closer inspection, we see the stochastic frequency trajectory of the adaptive mutation that structures the coalescent during the sweep. Here, the *a* and *bc* lineages recombine out of the sweep, and although the events occur in rapid succession, the remaining lineages do indeed coalesce pairwise. Prior to the sweep, neutral coalescence of the remaining lineages continues until a common ancestor is found.

Here we measure time pastward from sampling (*T*_0_ = 0) in units of 2*N* generations, i.e. on the coalescent time scale. We consider a single selective sweep of a *de novo* beneficial (and co-dominant) mutation with selection coefficient *s* that occurred at a discrete time point *T_a_* in the past. We define *T_a_* as the amount of time between fixation of the beneficial mutation and the time of sampling so that *T_a_* ≥ *T*_0_ = 0.

In the full evolutionary model, the beneficial mutation sweeps through the population following a stochastic frequency trajectory *X*[*t*] satisfying *X*[*T*] = 1 for *T* ≤ *T_a_* and for some 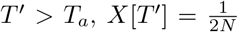 and *X*[*T* > *T*’] = 0. That is, the beneficial mutation arises as a single new mutation in a randomly-chosen background at time *T*’.

This frequency trajectory structures the coalescent process at linked neutral sites (Durrett and Schweinsberg, 2004). Coalescence occurs only between lineages which share the state at the selected site. Lineage pairs currently associated with the beneficial (conversely, ancestral) allele may coalesce at rate 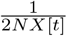 (respectively, 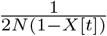), while any single such lineages may recombine out of (i.e. into, forwards in time) the sweep at rate *r*(1 – *X*[*t*]) per generation (respectively, *rX* [*t*]). Here, *r* is the rate of recombination between the selected and the neutral site.

Following the notations of Lohse et al. (2011), Hermisson and Pfaffelhuber (2008), and Barton et al. (2004), we label a sample of *n* lineages as a set {*a,b,c*…} and define the coalescence of the sample as a process that takes values in the set of partitions of {*a,b,c*…}. Each such partition is a set Ω = {Ω_1_,…,Ω_|Ω|_} such that 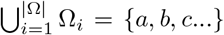 and Ω_*i*_ ⋂ Ω_*j*_ = 0 for *i* ≠ *j*. The process starts with the set of sampled lineages Ω = {{*a*}, {*b*}, {*c*}…} and ends when all lineages coalesce, Ω = {{*a, b, c*, …}}.

#### 2.1.1 Embedding sweeps in the Kingman coalescent, ϕ

For times *T* < *T_a_* i.e. before the selective sweep occurs, the ancestry of the sample Ω = Ω_*T*_ is described by the Kingman coalescent (Kingman, 1982). Any two distinct lineages 1 ≤ *i* < *j* ≤ |Ω| may coalesce with rate 1. In our setnotation, we represent this event by the removal of lineages *i* and *j* from the set Ω and replacing them with a single lineage representing their common ancestor: Ω → (Ω \ {Ω_*i*_, Ω_*j*_}) ∪ {∪_*i*_ ∪ Ω_*j*_}.

Following Lohse et al. (2011), we initially treat the sweep as a competing exponential process occurring at rate *δ* backward in time. This allows us to obtain through recursion the GF for the distribution of branch lengths in the genealogical history and, in a second inversion step, recover the GF parameterized by the discrete time when the beneficial mutation reaches fixation *T_a_*.

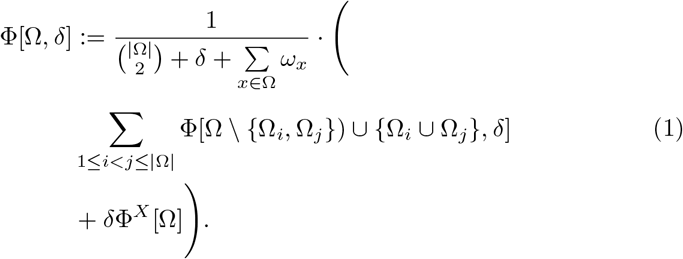

Here, ϕ^*X*^ represents the GF for the effect of the adaptive event (frequency trajectory of *X*) on the genealogy of our sample. In this paper, we focus on two different instantaneous sweep approximations: the star-like approximation and the instantaneous Yule approximation.

#### 2.1.2 The star-like approximation, ϕ*

In the star-like approximation, lineages independently recombine out of the sweep each with probability *P_e_* = 1 – *e^-α^* where 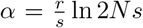 is a compound parameter for the relative distance from the sweep center, in which, *r* is the rate of recombination between the two sites, *N* is the (diploid) populaion size, and *s* is the heterozygous advantage of the beneficial mutation. The non-recombinant lineages that do not ’escape’ the sweep coalesce instantaneously to the origin of the beneficial mutation (Barton, 1998, 2000; Durrett and Schweinsberg, 2005). In the Kingman coalescent, only pairs of lineages may coalesce. Here, by contrast, coalescence may occur among any single subset of lineages of any size (including all or none). The possible subsets of Ω (formally, the power set of Ω, ℘(Ω) = {*S* ⊆ Ω}) represents all possible ways in which lineages may coalesce during the sweep. The effect is an instantaneous re-partitioning of the sampled lineages, and the recursion is defined as

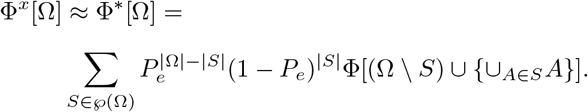

Note that ϕ on the right-hand-side no longer depends on δ, so that the sweep may only occur once in the genealogical history. Furthermore, ϕ* serves only to re-partition the sample. No length is added to the branches (no dummy variable terms *ω_x_*). We use the star-like approximation for analyses in the main text, and compare this to the more sophisticated approximation below.

#### 2.1.3 The instantaneous Yule approximation, ϕ**

Provided 2*N_e_s* is sufficiently large, the trajectory *X*[*t*] of a beneficial mutation under (strong) selection can be more closely approximated by considering a pure birth process with binary splits at rate 2*N_e_s* (Schweinsberg and Durrett, 2005). Forward in time, this Yule process describes the ancestry of all lineages descending from the beneficial mutation present at the end of the sweep (i.e. those with an infinite line of descent). Note that this process is stopped once there are 4*N_e_s* lineages, given that each lineage has a probability 2s of having an infinite line of descent. Genealogies under hitchhiking at neutral sites, at a distance *d* from the sweep site, can then be described by marking the Yule tree along its branches with recombination events occurring at rate *ρ* * *d*. A full recursion for this process is given in the appendix. Here we limit ourselves to using the sampling formula based on the Yule approximation (Etheridge et al., 2006; Bossert and Pfaffelhuber, 2013). Let Ω represent the set of lineages present at time *T_a_*, we can then define a labeled partition induced by the Yule process approximating the coalescent during the sweep. This partition consists of three families:

1. |*L*| = *l* late recombinant singletons: single lineages that have recombined away from the beneficial background.
2. Single family of early recombinants of size |*E*| = *e*: family of lineages recombining away after coalescing.
3. Single non-recombinant family of size |*N*| = |Ω| – |*L*| – |*E*|: family of lineages that are identical by descent to the founder of the sweep (along a distance of at least *d*).

Let Π** = {*S* = {*E,L,N*} : *E* ⋂ *L* = *E* ⋂ N = *L* ⋂ *N* = 0, *E* ∪ *L* ∪ *N* = Ω} be the set of unique partitions of Ω into classes *E, L*, and *N* defined as above. Using *P*[|*E*| = *e*, |*L*| = *l*] from Bossert and Pfaffelhuber (2013) and accounting for the 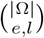 ways to uniquely partition the sample,

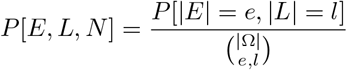

we obtain

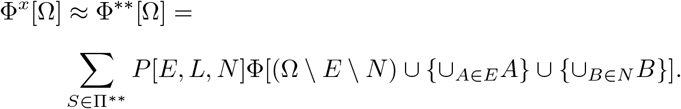

### 2.2 Simulations

The full model is implemented as a Wright-Fisher simulation using `SLiM3.3` (Haller and Messer, 2019) and `msprime` (Kelleher et al., 2016). Samples are extracted at a fixed number of generations after the sweep completes. Sequences are always 1Mb in length, with the site under selection in the center. We assume a population with *N_e_* = 10, 000, *r* = 1e^-7^ and *μ* = 1.25e^-7^ throughout, and simulate samples of varying size (*n* ∈ [4,12, 20]), sweep times (*Ta* ∈ [0.1, 0.5, 1.0, 2.0]) and two different strength of selection (*s* = 0.05 or 0.005). Simulation scripts and code for extracting genealogical information and the bSFS from the treesequences are available at https://github.com/GertjanBisschop/SweepsInTime.

### 2.3 Power Analysis

We assess the power to identify sweeps and the accuracy to infer sweep parameters (*T_a_* and *s*) using a composite likelihood (*CL*) scheme based on the blockwise site frequency spectrum (bSFS, see eq. (1) in Lohse et al. (2011)). Neutral variation on either side of a putative sweep target is summarized by the bSFS in *B* blocks of a fixed length *l* as a vector *k*. Blocks immediately to the right and left of the sweep target have an average distance *l*/2. Although we may sample a larger number of individuals *n*, analytic results for the bSFS are limited to smaller sample sizes. In the *CL* framework, we accommodate this by considering all possible subsamples of size *x* (we use *x* = 4 throughout). Let *P*[*k_ij_*] be the probability of observing a block-wise mutation configuration *k* at distance *i* * *l* –1/2 in subsample j, 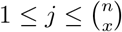. Summing over all 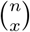 subsamples of *n*, we can define the following composite likelihood for the sweep model,

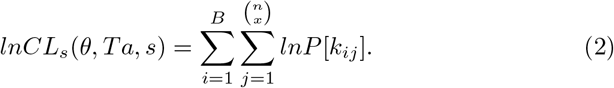

Given an analogous likelihood under neutrality *lnCL*_0_(*θ*) the support for a sweep (at time *T_a_* and of strength *s*) can be measured as:

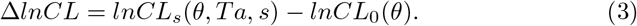

We fit both models to 10,000 simulated replicates with a beneficial mutation as well as to 10, 000 neutral simulations. To measure power we construct ROC curves: Δ*lnCL* values for *true* (hard sweep) and *false* (neutral) replicates are jointly ranked in descending order, after which for each element both the fraction of false and true positives are determined. Note that we do not perform a sweep scan but rather assume that the position of the selective target is known.

Rather than evaluating all equations for each combination of parameter values (*θ*, *Ta*, *s*, *N_e_*, *r*), we construct an interpolation function (third degree polynomial) in *Mathematica* (version 12) for each *P*[*k_ij_*] to reduce computation time significantly.

For each replicate, inference consists of two steps: we first estimate *θ*, using all blocks that are sufficiently far away from the sweep site (*α* > 12). We then obtain joint estimates of *Ta* and *s* conditional on *θ*.

Parameter optimization was conducted on a grid (*θ*, *Ta*, *s*) allowing us to pre-compute all bSFS configurations, and run the optimization for all simulation replicates on a laptop.

## 3 Results

### 3.1 Time erodes the footprint of adaptive evolution

In this section, we examine how the expected footprint of adaptive evolution is affected by *T_a_*, the time since the selective sweep. We first show results for pairwise genetic diversity (*n* = 2) and then extend this to the site-frequency spectrum (*n* = 10). A detailed analysis as well as an illustration of our approach is provided in S1 Notebook.

#### 3.1.1 Pairwise Genetic Diversity

For illustration, we first derive the GF for the distribution of branch lengths in a sample of two (haploid) lineages. In this case, the two branches a and b are equivalent, and their sum is twice the time to the most recent common ancestor (*t_mrca_*). By substituting *ω_a_* + *ω_b_* → *ω_mrca_* = *ω*, we obtain the GF for *t_mrca_*. Under the neutral model, *φ*_2_ is simply the Laplace transform of an exponentially distributed random variable with mean 1

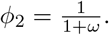

For the selective sweep model, the recursion for the GF with parameter *δ* is:

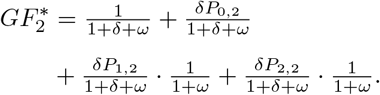

The summed terms represent the possible sequences of events in the genealogical history of the sample. The first term corresponds to neutral coalescence before the sweep, the second to coalescence during the sweep. In the remaining terms, one or both lineages escape the sweep and subsequently coalesce under the standard neutral coalescent. Inverting 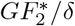 with respect to *δ* gives the GF for the distribution of branch lengths as a function of the time since the sweep *T_a_*:

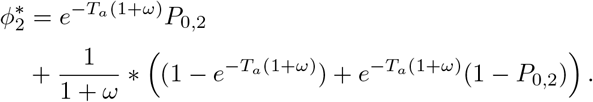

Setting *ω* → 0, we obtain the probability for each genealogical history (Lohse et al., 2011). Neutral coalescence occurs before the sweep with probability 1 – *e^−T_a_^*. Given that it does not (with prob. *e^-T_a_^*), coalescence may happen during the sweep with probability *P*_0,2_ or neutrally after the sweep with probability (1 – *P*_0,2_). The expected time to the most recent common ancestor 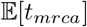 is obtained by taking minus the derivative of the GF with respect to *ω* and then taking the limit as *ω* → 0 (Lohse et al., 2011). In the neutral case, 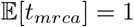, and for the sweep scenario, substituting *P*_0,2_ = *e*^-2*α*^, 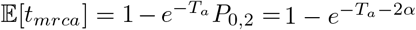.

When *T_a_* = 0, i.e. the population is sampled at the time of fixation, we recover the classic valley of diversity caused by a selective sweep (Maynard Smith and Haigh, 1974; Kaplan et al., 1989a) (Fig. 2, A). By comparison, older sweeps have a reduced effect on 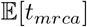. Forwards in time, this amounts to the recovery of genetic diversity that was lost due to hitchhiking in the selective sweep. From a coalescent viewpoint, the genealogy is unaffected by selection if coalescence occurs before the sweep.

**Figure 2:**
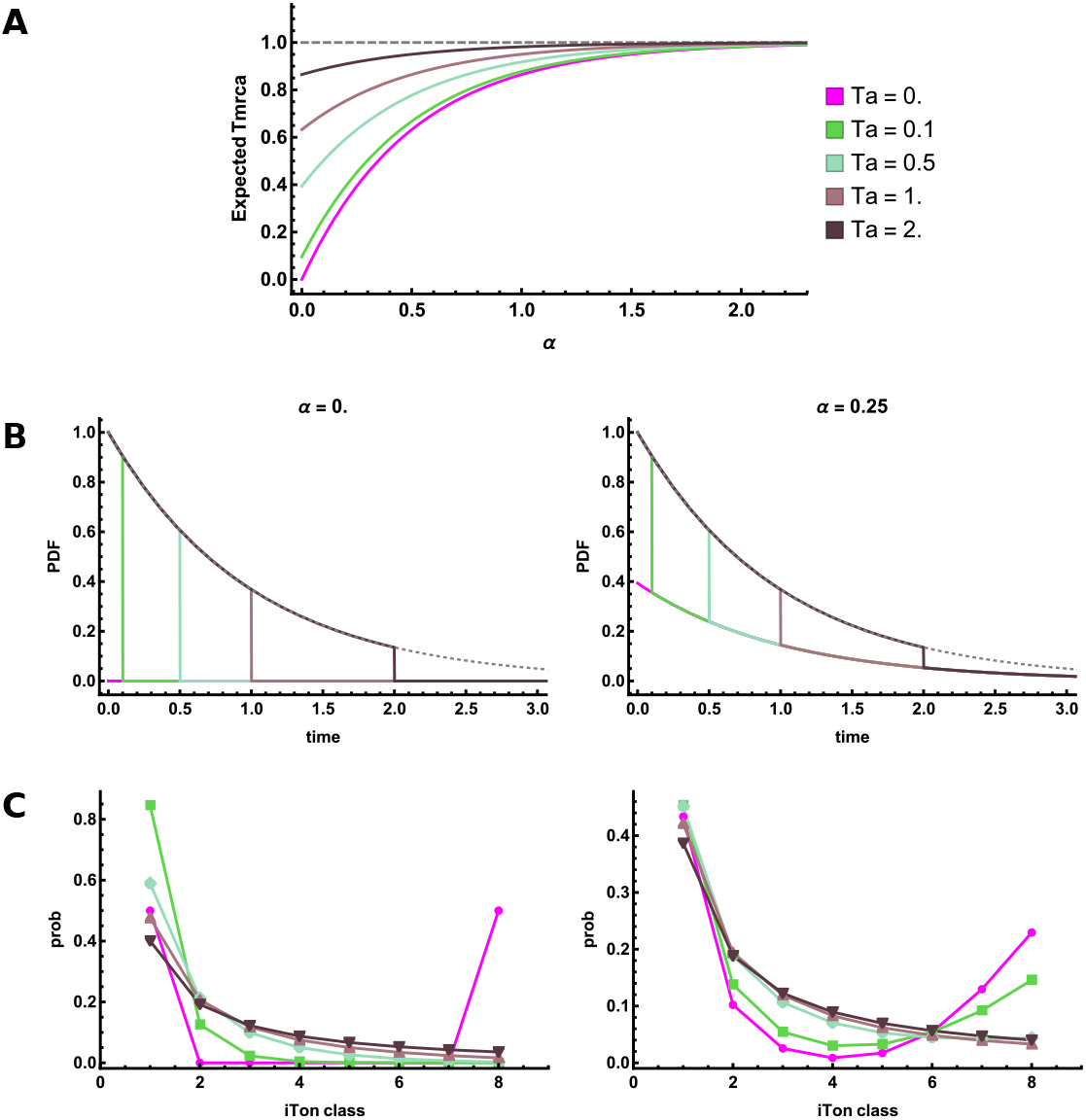
A: the extent to which expected genetic diversity is reduced around a sweep and its recovery with distance from the sweep centre 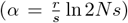 depends on the time since the sweep *T_a_*. B: the (probability density) distribution of the time to the most recent common ancestor at the sweep centre, *α* = 0 and at distance *α* = 0.25. Panel C: the site frequency spectrum for a sample of n = 9 individuals.

#### 3.1.2 Distribution of *t_mrca_*

The effect old sweeps have on the genealogy can be seen more clearly in the full distribution of *t_mrca_:* under the neutral model, *t_mrca_* is exponentially distributed with rate 1. The probability density (PDF) and cumulative density functions (CDF) are therefore *f* [*t*] = *e^-t^* and *F*[*t*] = 1 – *e^-t^* respectively. Under the selection model, we obtain the PDF by inverting the GF with respect to *ω* (Lohse et al., 2011). We may integrate the PDF with respect to t to obtain the CDF or alternatively, we may divide the GF by *ω* and then take the inverse Laplace transform. For this model, we obtain

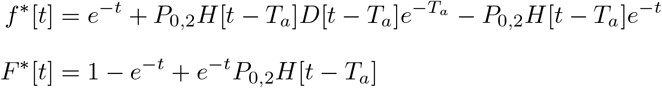

where *D*[*t*] and *H*[*t*] denote the Dirac delta and Heaviside step function respectively. As expected, for times *t* < *T_a_*, the PDF matches the neutral case, *f* *[*t*] = *f*[*t*], since the sweep cannot affect the genealogy during that period (Fig. 2 B). Since we assume that the sweep induces an instantaneous coalescent event, there is a point mass of size *e*^-*T_a_*^ *P*_0,2_ = *e*^-*T_a_*-2*α*^ at *t* = *T_a_*. Indeed, at the sweep center, all coalescence occurs before or during the sweep *t* ≤ *T_a_*. At greater distances from the sweep center, the point mass diminishes in size. For *t* > *T_a_*, only lineages that escaped the sweep may subsequently coalesce, and they do so neutrally. Thus, the probability density matches the neutral case scaled by the probability that one or both lineages escape, *f*\*[t] = *e^-t^*(1 – *P*_0,2_) = *f*[*t*](1 – *P*_0,2_). Indeed, Fig. 2 shows that, although the location of the discontinuity shifts as the time since the sweep increases, the probability density for *t* > *T_a_* is determined only by the distance from the sweep center, 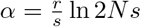.

#### 3.1.3 The site frequency spectrum

For moderate sample sizes, we can obtain the expected site frequency spectrum (SFS) as a function of both the distance from the sweep center *α* and the time since the sweep. Distinguishing branches by the number of descendants, e.g., *ω_a,b_* → *ω*_2_, the set of *ω_i_*, 1 ≤ *i* ≤ *n* – 1 corresponds to the length of the branches with *i* descendants among the sample (i-Tons). The expected *marginal* lengths of i-Ton branches 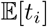 can be obtained by differentiating the GF with respect to *ω_i_*, analogous to 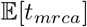 described above for *n* = 2. Normalizing by 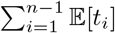 yields the expected frequency of mutations belonging to each i-Ton class. Fig 2 C, shows the SFS for a sample of *n* = 9, at different distances from the sweep center and for increasingly old sweeps. As for the expected pairwise genetic diversity, the effect of older sweeps on the SFS is dampened. However, the relative effect differs between i-Ton classes and depends on both *T_a_* and *α*. At the sweep center, *α* = 0, we observe a prominent excess in the proportion of singleton lineages. In contrast, outside the sweep center, *α* = 0.25, we see an excess of both intermediate and high frequency polymorphisms as the age of the sweep increases.

### 3.2 Beyond the mean – leveraging joint branch length information

Pairwise and/or average measures of sequence variation such as the SFS are drastic summaries of sequence variation. In order to fully capture the footprint of selective sweeps on linked neutral sequence variation, we would ideally like to compute the probability of haplotypic variation flanking a selective target. Unfortunately, this requires including recombination (including breakpoint locations) explicitly in the GF recursion which quickly becomes intractable. In the following, we focus on blocks of non-recombining sequence and consider the effect of sweeps on three quantities of interest: the probability of genealogical topologies, the marginal distribution of i-Ton branches and, following Lohse et al. (2016), the bSFS, the blockwise configuration of i-Ton counts.

#### 3.2.1 The probability of genealogical topologies

The probability of seeing any particular topology can be found by taking the limit of *ω* corresponding to branches that are incompatible with it (and evaluating all other *ω* at zero). Under *φ\** and for *n* = 4, this results in five different topologies, three of which are induced by multiple-mergers. For the sake of simplicity, we can distinguish between three topology classes defined by the root node: genealogies with a symmetric or asymmetric bi-partition at the root and genealogies without any bi-partition (*P_star_*) (see Fig. 3).

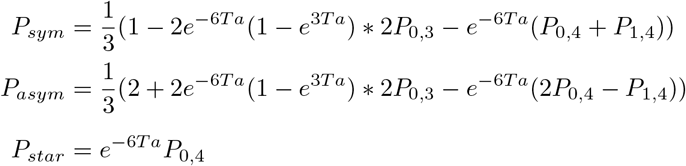

**Figure 3:**
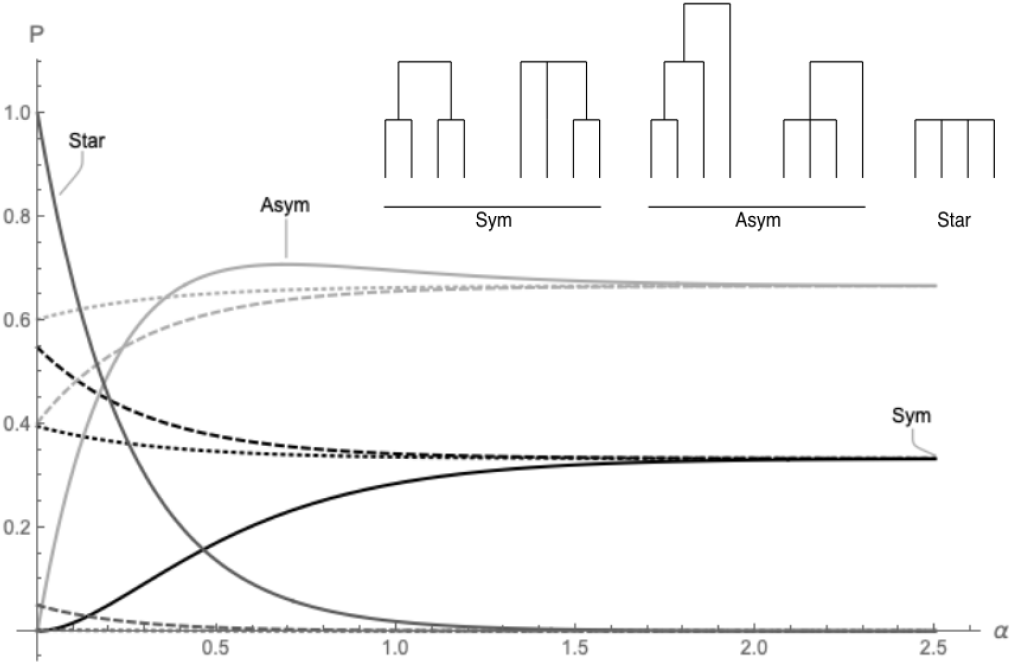
Probability of genealogical topologies. The probability of a genealogy with an asymmetric (light gray), symmetric (black) or star-shaped (dark gray) root-node is shown for *T_a_* = 0.0 (full), 0.5 (dashed), 1. 0 (dotted), with increasing distance from the sweep centre (left to right).

Dissecting the three terms in *P_sym_* and *P_asym_,* the first term represents the probability of seeing (a)symmetric trees under the standard neutral coalescent. Only multiple-mergers of three (second term) or four (third term) lineages will affect this probability.

#### 3.2.2 The marginal distribution of i-Ton branches

The marginal distribution (PDF) of i-Ton branches, i.e., the genealogical branches underlying the SFS (Fig. 2 C), can be obtained by inverting the GF with respect to ω_*i*_. The resulting expressions are cumbersome. They are obtained using the machinery in S1 Notebook, where we also provide them explicitly for a sample size of four, which we investigate below.

For the case of *n* = 4 and assuming neutrality, only two topologies are possible: the first coalescent event always generates a doubleton lineage, while the second may either generate a second doubleton lineage, resulting in a symmetric topology with probability *P_sym_* = 1/3, or a tripleton lineage, resulting in an asymmetric topology with probability *P_asym_*2/3. Thus, the marginal PDF of tripleton branches *f* [*t*_3_] contains a point-mass at *t_3_* = 0 of size 1/3, while the PDF for *t*_3_ and *t*_2_ contains no discontinuities (Fig. S1 A).

In contrast, in the vicinity of a selective sweep, we observe multiple discontinuities in all three marginal PDFs (Fig. 4). However, given that each class of i-Ton branches consists of multiple genealogical branches, these are more intricate than for the pairwise pairwise case (*f* [*t*_2_] above). For example, there are three discontinuities in the PDF of singleton branches: removable discontinuities at *t*_1_ = *T_a_* and *t*_1_ = 2*T_a_* and a jump discontinuity at *t*_1_ = 4*T_a_* with a point mass of size *e*^-6*T_a_*^*P*_0,4_, corresponding to the case where the first event is the simultaneous coalescence of all lineages during the sweep. The PDF of *t*_2_ contains two removable discontinuities, one at *t*_2_ = *T_a_* and one at *t*_2_ = 2*T_a_* and one point mass at *t*_2_ =0 of size *e*^-6*T_a_*^ (*P*_0,4_ + *P*_1,4_). This point-mass is again caused the by multiple-merger coalescence events that give rise to a topology devoid of doubleton branches. For *t*_3_, we observed a removable discontinuity at *t*_3_ = *T_a_* and, as for the neutral case, a point-mass when *t*_3_ =0. The size of this point mass, given below, equals *P_sym_* + *P_star_* (see above), and so is no longer a simple function of the *P_i,k_* under the star-like approximation, as *P_sym_* may result from different combinations of neutral coalescence events and sweep induced coalescence, whereas *P_star_* = *P*_0,4_ corresponds to the probability that all lineages coalesce in a sweep-induced multiple merger event:

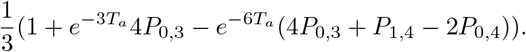

**Figure 4:**
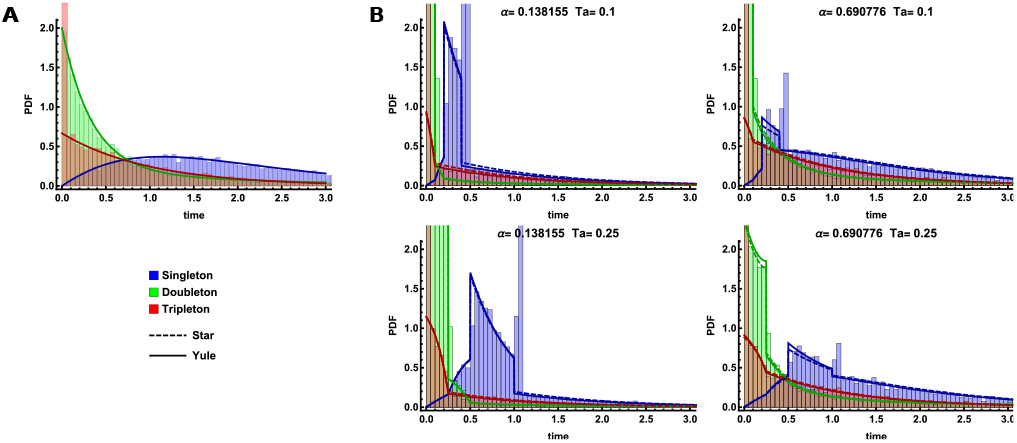
Marginal i-Ton branch length distributions for *n* = 4. Analytic predictions under the neutral model (A) and the approximate selection models (B) are compared to the corresponding distribution obtained from 10, 000 simulation replicates overlaid as a histogram. The Yule approximation is indicated by solid lines while the dashed lines indicate the star-like approximation. Singleton, doubleton and tripleton branch lengths correspond to the colors blue, green and red respectively. The top row shows two distances from the sweep center *α* ≈ {0.14,0.069} and *T_a_* = 0.1. Analogous results for an older sweep *T_a_* = 0.25 are shown in the bottom row. Time is measured in units of 2N generations. Here, *N* = 10,000, *s* = 0.05, and *r* = 1*e* – 7.

More generally, for a sample of size n, the discontinuities present in each i-Ton branch type are determined by the total number of each such branch present during the inter-event times of the coalescent process during which the selective sweep occurs. For singleton branch types, there are always *n* 1-Ton branches present initially, and the first coalescent event reduces this to *n* – 2. There exists one topology in which the number of singletons diminishes by one each subsequent step. Therefore, there are singleton discontinuities at *t* = {(*n*)*T_a_*, (*n* – 2)*T_a_*, (*n* – 3)*T_a_*,…, *T_a_*}, for a total number of *n* – 1. For *i* > 1, there is always a discontinuity and point mass at *t* = 0 due to the possibility that the first event is coalescence of all lineages during the sweep. The possible multiplicity of the (*i* > 1)-Ton classes is determined by the ways to decompose *n* into smaller-valued integers and thus the number of discontinuities for *i* > 1 is [n/ij + 1.

Finally, we note that the star-like approximation used for the analysis provides relatively accurate predictions in this case. In comparison to simulations, the accuracy improves only slightly using the Yule approximation (Fig. S1).

#### 3.2.3 The blockwise site frequency spectrum

Above, we used the GF to obtain the SFS by deriving the expected length of i-Ton branches. An alternative and less drastic summary of sequence variation is the bSFS, the vector of SFS counts in short blocks (Bunnefeld et al., 2015). Assuming no recombination within blocks, the bSFS can be obtained from the GF by taking successive derivatives wrt to the ω_*i*_ (see eq. (1) in Lohse et al. (2011) for details). Note that for the minimal sample size of *n* = 2 the bSFS reduces to the distribution of pairwise differences (Wang and Hey, 2010; Wilkinson-Herbots, 2008). To be able to leverage topology information, we will focus on (sub)samples of *n* = 4. In this case, the bSFS is a vector of counts for three i-Ton types 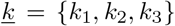 where *k_i_* ∈ {0,1…, *k_max_*}. Note that bSFS configurations with *k*_3_ > 0 imply an underlying genealogy with tripleton branches, i.e. 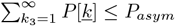

Comparing the analytic expectation for the bSFS 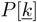 to simulations (Figure 5) highlights both the accuracy of the star-like approximation (for *n* = 4) and the robustness of the bSFS to intra-block recombination provided blocks are short (here a recombination rate of *r* = 10^-7^, *l* = 100 bases). If we restrict the bSFS to a maximum of *k_max_* = 2 mutations per i-Ton type, we distinguish (*k_max_* + 2)^3^ = 64 unique bSFS configurations (given that both the absence of a particular mutation type *k_i_* = 0 and seeing > *k_max_* mutations define bSFS configurations).

**Figure 5:**
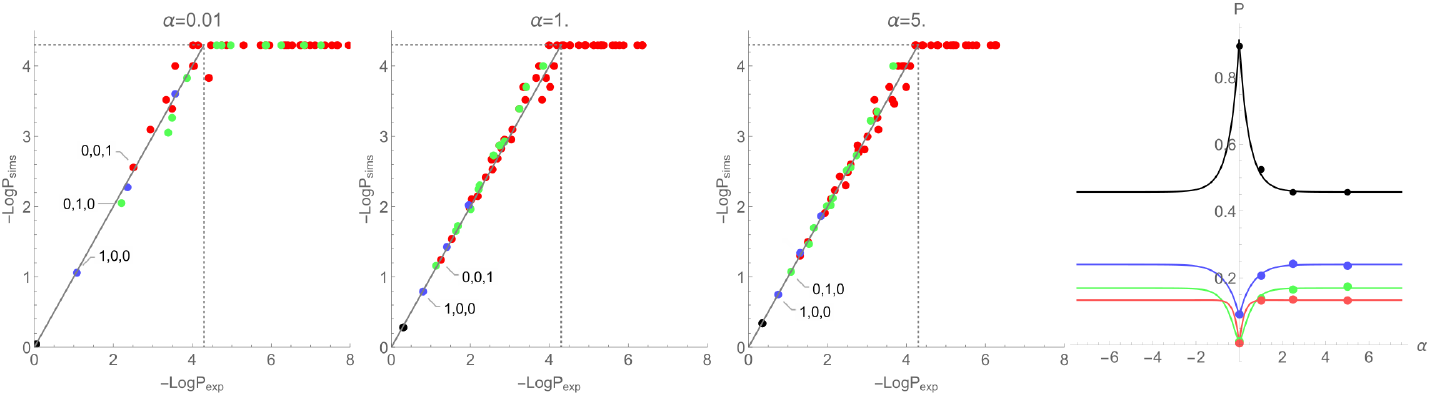
The blockwise site frequency spectrum *n* = 4,*T_a_* = 0.1. The expected probabilities of bSFS configurations given by *ϕ*\* (logscale) against their observed frequencies in 10,000 simulation replicates. Each dot corresponds to a unique bSFS configuration. Counts left and right of the selected site are added together. Each dot in the scatter plot represents a unique bSFS-configuration, counting the number of (singletons, doubletons, tripletons). Red: (_, _, *k*), green: (_, *k*, 0), blue: (*k*, 0,0), black: (0,0,0) with *k* ≤ 1, and _ any integer. The dotted line marks the minimal detectable frequency for the simulations. The rightmost figure shows the total probability of observing blocks within each of these categories (sweepsite at *α* = 0.0).

### 3.3 Power to infer historical sweeps

We can use the analytic result for the bSFS obtained above using the GF to jointly estimate the sweep time and the strength of selection in a composite like-lihood framework (summing *lnL* across both blocks and subsamples of *n* = 4, see Methods). In the following, we quantify the power (and bias) of characterising sweeps using the star-like approximation (*φ*^*^) and test to what extent the instantaneous Yule approximation (*ϕ*\*\*) improves these estimates.

With strong selection (*s* = 0.05), the power to infer *T_a_* and *α* is high, even for fairly old sweeps (*T_a_* = 1.0), especially with *n* ≥ 12 (see Fig. 6). Even for small sample sizes, we get decent estimates of the sweep parameters (see Fig. 7). Increasing sample size (for a fixed subsample size of *n* = 4) will reducing the mutational sampling noise, but only increases power to estimate parameters to a limited extent (see Fig. S2). The power to correctly infer sweep parameters decreases with increasing age of the sweep (*T_a_*). This is unsurprising, given that the number of lineages that enter the sweep, and hence the information about it, declines with increasing *T_a_*.

**Figure 6:**
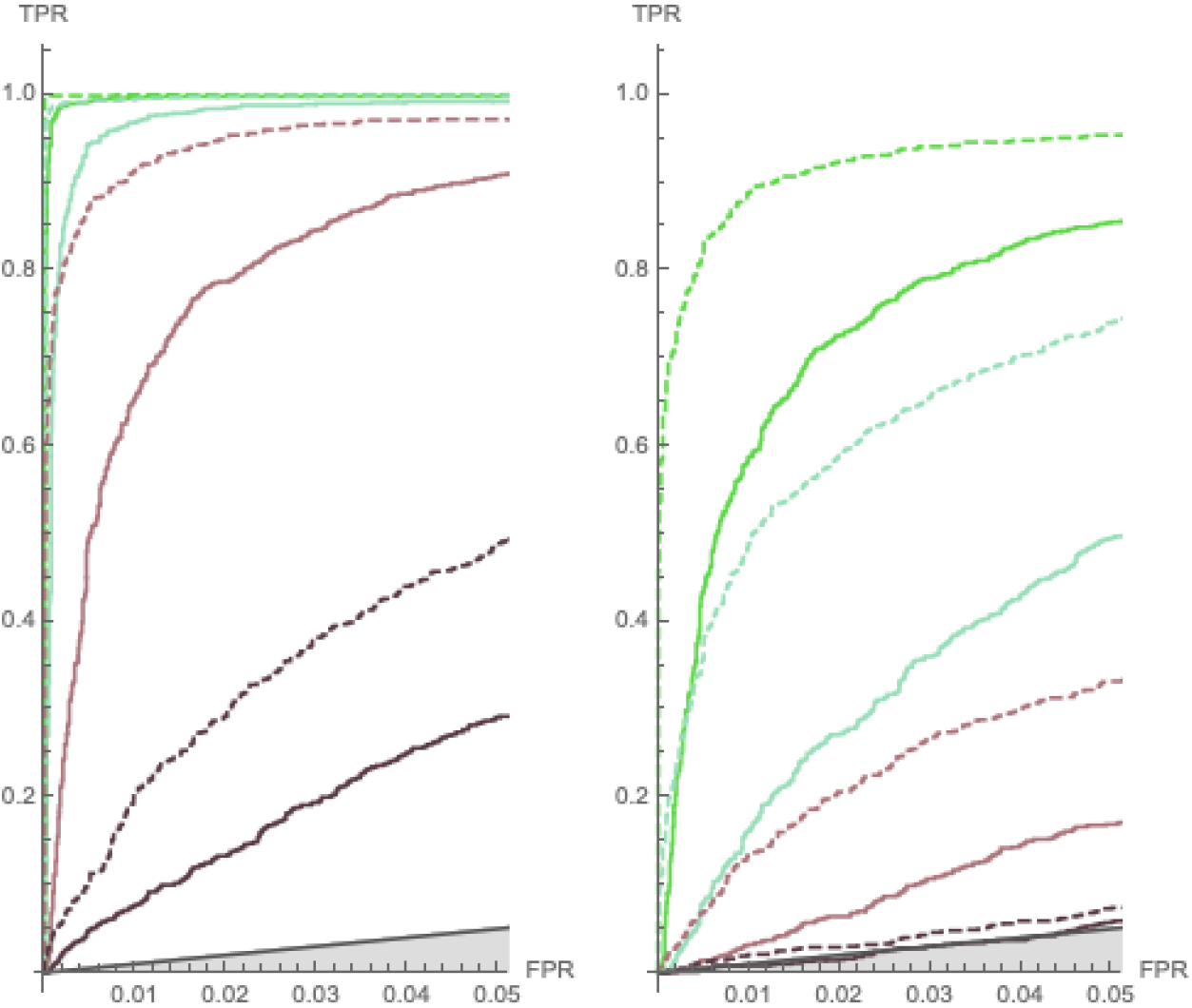
ROC curve. Plotting the rate of true positives against the rate of false negatives shows how much power we have to distinguish genomic regions that underwent a hard sweep from neutral replicates. As expected, power depends on the time since the sweep (*T_a_* = 0.1 (green), 0.5 (lighter green), 1.0 (light brown), 2.0 (dark brown)), the strength of selection (left *s* = 0.05, right *s* = 0.005) and sample size *n* = 4 (full line), 12 (dashed).

**Figure 7:**
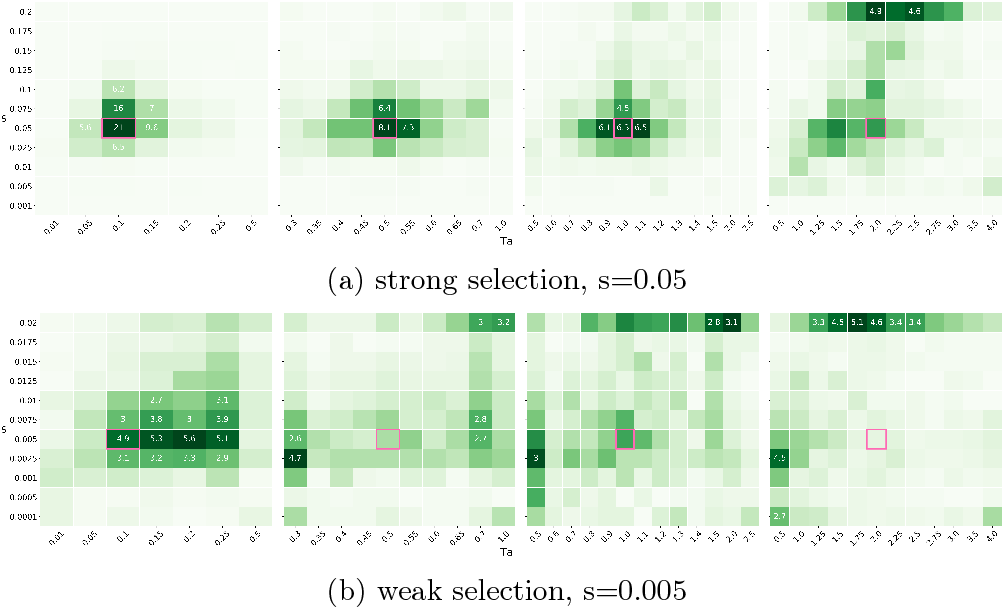
Heatmaps. Results of the gridded optimization. Panels represent different sweep ages (*T_a_* = 0:1; 0:5; 1:0; 2:0 from left to right). Numbers show the percentage of replicates associated with a particular parameter combination. The true parameter combination is indicated by a pink square.

When selection is weak (*s* = 0.005), power drops off quickly for sweeps older then *T_a_* = 0.5, especially for small samples (*n* = 4). The heatmap reveals that sweep parameters become non-identifiable in this weak selection limit independent of sample size (see Figs. 7 and S3). By contrasting ROC curves between a model with *T_a_* = 0 and a model where *T_a_* is free to vary, we can assess how much better the data fits a historic sweep (see Fig. S4). As expected, forcing *T_a_* = 0 (a standard assumption in sweep scans), works well for recent sweeps but breaks down for older ones, i.e. power drops off in the same way previously reported by SFS-based methods (Racimo et al., 2014; Setter et al., 2020). However, when including the sweep time as a parameter, historic sweeps become detectable with high power as long as they are strong.

Comparing the heatmaps and ROC curves under the star-like and the instantaneous Yule approximation (see Figs. S5, S6, and S7) reveals that there is very little difference between the two approximations in terms of accuracy and power. Root mean square errors for the estimates are nearly identical across a range of *T_a_* estimates. We also find that old sweeps are similarly non-identifiable under both approximations when selection is weak. Thus the power to infer selection is inherently limited in this case.

### 3.4 Out of Equilibrium - Adaptive Introgression

So far, we have assumed that there is no population structure and that, with the exception of a focal sweep, the population is at equilibrium: the adaptive mutation arises *de novo* in an otherwise neutral panmictic population of constant size. Ideally, we would like to infer a history of selection in the context of a known demographic model, e.g. to identify sweeps associated with range expansions, domestication, or speciation events. The GF framework with selection allows for such joint inference. As an initial first step, and to demonstrate the exibility of the GF framework, we analyzed the model of adaptive introgression after secondary contact of Setter et al. (2020). We highlight the main results and refer to the detailed analysis in S2 Notebook. In this model, we assume that a population split occurs at some time *T_s_* in the past and results in two fully-isolated populations. We further assume that an adaptive mutation fixes in one population (the donor), and, through secondary contact at time *T_a_*, a single haplotype carrying a beneficial mutation migrates from the donor to the recipient population and subsequently sweeps to fixation. We assume through out that the time interval between the divergence of the donor population and the adaptive introgression event *T_d_* = *T_s_* … *T_a_* is fixed and explore the effect of the age of the introgression sweep (i.e. the time *since* the sweep occurred *T_a_*) on the genealogies at nearby neutral sites.

As for the classic sweep scenario, we find an increasingly dampened signal of adaptation as *T_a_*, the time since the sweep, increases, and unless the sweep is relatively recent, simple measures of genetic diversity confound *T_a_* and *T_s_,* the time since the populations split. However, these time parameters have clearly distinguishable effects on the marginal branch length distributions (Fig. S8 and Fig. S9). As for the classic model, the burst of coalescence observed for singleton and doubleton branches can only be generated by selection. All singleton branches coalesce in a single burst if the first event is the selective sweep and none of the lineages escape. Similarly, for doubletons, a point mass at *t* = 0 results from a multiple merger of only three individuals during the sweep. The size of these point masses depends only on *T_a_*, and the divergence from the donor *T_d_* shifts the distribution pastward. However, for tripleton lineages, the size of the pointmass at *t* = 0 also depends on the time since the split, and this is due to the structure imposed on the coalescent by the strict isolation model. For example, suppose the sweep is the first event in the local genealogy, and three out of four lineages escape. If the subsequent time to the population split (*T_d_*) is large, we expect all three of the escaped lineages to coalesce before finding a common ancestor with the donor lineage. In other words, for this sequence of events, the probability of observing tripleton branches is higher than under the classic sweep model due to the divergence time.

Although we saw little difference in the distribution of i-Ton branches between the star-like and the instantaneous Yule approximation for a classic hard sweep, in the case of introgression sweeps the Yule approximation does provide noticeably more accurate prediction than the star-like approximation.

## 4 Discussion

We have shown how the effect of selective sweeps on nearby genealogies can be modelled analytically using the recursion for the generating function for the genealogical histories (Lohse et al., 2011). Much like a population bottleneck which can also be approximated as a multiple merger event (Bunnefeld et al., 2015), a selective sweep can be viewed as a discrete event that affects the genealogical history of a sample of nearby neutral lineages (Kaplan et al., 1989b). However, unlike population bottlenecks, selective sweeps have a local effect on neutral variation in the genome (Galtier et al., 2000) and lead to topologically unbalanced genealogies, and so are distinguishable.

While it is straightforward to recover previous analytic results for the expected loss of pairwise genetic diversity around sweep targets (Maynard Smith and Haigh, 1974; Kaplan et al., 1989b) and the SFS using the GF framework, our motivation was to extend analysis beyond expected coalescent times and pairwise samples. What we gain by embedding selective sweep approximations in the GF framework is a complete analytic description of the effects of genetic hitchhiking on the distribution of genealogies. Crucially, the strength and age of selective sweeps distort genealogies at nearby neutral sites in distinct ways. While these two aspects of past selective events are hard to disentangle from the expected reduction in genetic diversity, we show that they can be jointly estimated using richer summaries of sequence variation which capture information contained in the distribution of genealogies. Specifically, we show that for a single strong selective sweep, the bSFS has reasonable power to jointly infer both parameters even for a sample of *n* = 4 lineages. Being able to maximise the information contained in small samples not only provides an obvious avenue for composite likelihood inference, but also increases the power of comparative population genetic analyses, which are still limited by the lack of large re-sequence data sets for most taxa. While we have limited ourselves to exploring the bSFS and the problem of inferring selection parameters around known sweep targets as a proof of principle, it should be possible to extend this framework to build genome scans for historic sweeps that occurred at a specific time of interest (*T_a_*).

### 4.1 Model extensions and limitations

#### 4.1.1 Star-like vs Yule

Throughout this paper we have focused on two sweep approximations. While the instantaneous Yule approximation is a more accurate description of a hard sweep than the star-like approximation, we find very little difference in terms of power and accuracy between both sampling formulas in the case of a classic hard sweep. However, it may be unsurprising that ignoring the possibility of a family of early recombining lineages has little impact given that the (sub)sample size we considered is small (Pfaffelhuber et al., 2006).

For the more complex adaptive introgression scenario, we do observe a better fit for the length distribution of i-Ton branches under the Yule approximation. This suggests that it is worth considering the difference in accuracy between both approaches when implementing more complex scenarios of selection. The (instantaneous) Yule approximation does come at a computational cost. Firstly, the number of possible sampling configurations resulting from the sweep is 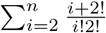 versus 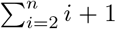 under the Yule and star-like approximation respectively. Secondly, the Yule approximation requires the calculation of a random variable, the computation time of which is proportional to 4 * *N* * *s*.

#### 4.1.2 Different types of selection

Although there has been much interest in differentiating the signatures of soft and hard sweeps (Hejase et al., 2020b), previous analytic work has shown that old hard sweeps are difficult to distinguish from soft sweeps given that both cause a partial reduction in genetic diversity (Hermisson and Pennings, 2005; Pennings and Hermisson, 2006b,a). While recent soft sweeps can be distinguished by conspicuous patterns in haplotype data (Ferrer-Admetlla et al., 2014), these associations break down relatively quickly, and an old soft sweep may be indistinguishable from a (slightly older) hard sweep (Zheng and Wiehe, 2019; Schrider et al., 2015). Despite this, machine-learning methods appear capable of classifying different histories of selection (Hejase et al., 2020b). By incorporating models of soft selective sweeps (Hermisson and Pfaffelhuber, 2008) into the GF framework, it should be possible to identify the characteristics signatures of these selective processes in the branch length distributions and/or gene tree topologies.

We focus on the effects of a single hard sweep. An alternative is to capture the aggregate effects of positive selection on patterns of neutral diversity throughout the genome (Booker et al., 2017; Juric et al., 2016). While the signal of any particular sweep is inherently limited (given the stochasticity of both the coalescent and the trajectories of selected alleles), one would expect much more information about positive selection when aggregating signatures across the genome. Given that our starting point has been to assume a model in which the waiting time to a sweep is exponentially distributed (see eq. 1). While, in the context of a single sweep (the scenario we have considered here) this was done purely for mathematical convenience, it also yields the recursion for the GF under a model of recurrent sweeps.

#### 4.1.3 Joint inference of demographic history and selection

The majority of theoretical results for selective sweeps to date have assumed that there is no population structure and that, with the exception of a focal sweep, the population is at equilibrium: the adaptive mutation arises *de novo* in an otherwise neutral panmictic population of constant size. In reality, of course, natural populations are not at equilibrium (Brandvain and Wright, 2016) and it has remained extremely challenging to jointly infer past demography and selective events (LI et al., 2012). The most successful approaches to date extend the approximate diffusion model of Kimura (1955) to describe the population-level allele frequency spectrum under non-equilibrium dynamics. However, solving the diffusion equation can be difficult. Zivkovic and Stephan (2011) obtain analytic results for historically varying population sizes, but in combination with positive selection, only numeric solutions are possible (Williamson et al., 2005) except for very simplistic demographic histories (Evans et al., 2007). Crucially, these predictions are primarily used to infer the effects of *direct* selection by comparing allele frequency spectra among different classes of mutations (e.g. coding vs. non-coding). While this approach can provide demographically explicit predictions for the background SFS in sweep-scanning methods (Pavlidis et al., 2013; Johri et al., 2020), results to-date are again limited to the SFS and to very recent sweeps (*T_a_* = 0).

As an initial first step, and to demonstrate the flexibility of the GF frame-work, we analyzed the model of adaptive introgression after secondary contact of Setter et al. (2020). We obtained predictions for measures of genetic diversity and marginal branch lengths as functions of both the strength of the sweep *α* and the time between successive demographic events, *T_a_,* the time to introgression, and *T_s_*, the divergence of the donor at the time of introgression. While these inter-event times are confounded in measures of pairwise genetic diversity, they are clearly distinguished from richer summaries of the data such as the marginal branch length distribution. Importantly and perhaps more subtly, the GF method provides further improvement: it fully accounts for the sorting of lineages and is accurate even if the donor population only recently diverged. Our results suggest that methods based on the GF are capable of inferring complicated histories of both demography and selection.

### 4.2 Towards more powerful inference of selection

The motivation for our analytic work is to improve the ability to make inferences about selection. While we have explored one approach, a composite likelihood framework based on the bSFS of sub-samples, in some detail, there are several other promising avenues for developing inference.

Our results for the effect of sweeps on genealogical branches may prove to be powerful in the context of recent methods that infer the ARG and/or tree sequences (with or without branch length information) from phased data such as ARGweaver (Rasmussen et al., 2014), and RENT+ (Mirzaei and Wu, 2017), tsinfer (Kelleher et al., 2019), and RELATE (Speidel et al., 2019). In principle, our GF framework allows to connect a sequence of marginal trees inferred by these methods to explicit models of population structure and past selection.

One direction of further research could build on the fact that tree sequences returned by tsinfer only contain topology information. It will be interesting to ask how much information is contained in the distribution of topologies, allowing us to extend the GF to the larger sample sizes required by these methods. Several summary statistics have been developed to diagnose the effect of sweeps on genealogical topologies (Li and Wiehe, 2013; Yang et al., 2018). This research is motivated by the fact that statistics like root imbalance are invariant to population size changes. But, as far as as we are aware, results for the effect of sweeps on the distribution of topologies are lacking and could be used to improve sweep scans. For example, the probability of asymmetric topology (i.e. a bi-partition of {3,1} in a sample of *n* = 4) follows a non-monotonic pattern around sweep targets. Analogous signals have been exploited to distinguish adaptive introgression sweeps from classic sweeps (Setter et al., 2020).

A final approach would be to compute the joint probabilities of the mutational configuration/branch lengths of a tree and its span. Leaving out the mutational information used to infer the tree sequences, inference would be based directly on the distribution of marginal genealogies, including the distribution of coalescence times (Weissman and Hallatschek, 2017). While a full model of recombination, i.e. allowing for an arbitrary number of recombination breakpoints in a sequence seems infeasible, it should be possible to condition the GF on there being no recombination in a stretch of sequence of a given length. Abandoning the idea of non-recombining blocks of a fixed length would thus allow us to incorporate LD information in the sweep inference. Although the direct inspection of the marginal trees that represent the genealogical history of a sample is an exciting prospect, we still require the statistical tools to exploit the information they contain about the evolutionary process efficiently.

## Acknowledgements

This work was supported by an ERC starting grant (ModelGenomLand, 757648). KL is also supported by a fellowship from the Natural Environment Research Council (NERC, NE/L011522/1). We thank Brian Charlesworth and Matthew Hartfield for insightful comments on an earlier draft.

## 5 Supporting Information

### SI files

#### S1 Notebook

Machinery and analysis of marginal branch lengths for classic hard sweep scenario.

#### S2 Notebook

Machinery and analysis of marginal branch lengths for adaptive introgression scenario.

### SI figures

**Figure S1:**
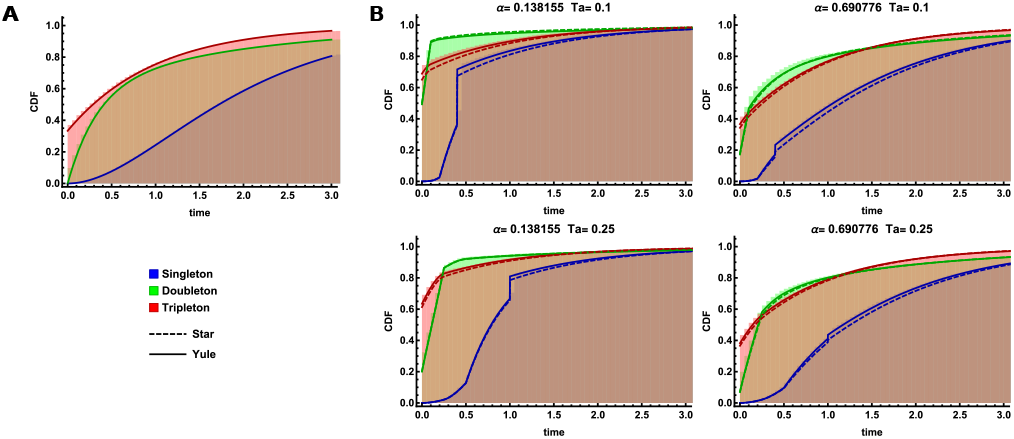
CDF for marginal branch lengths. Companion to Fig. 4 of the main text.

**Figure S2:**
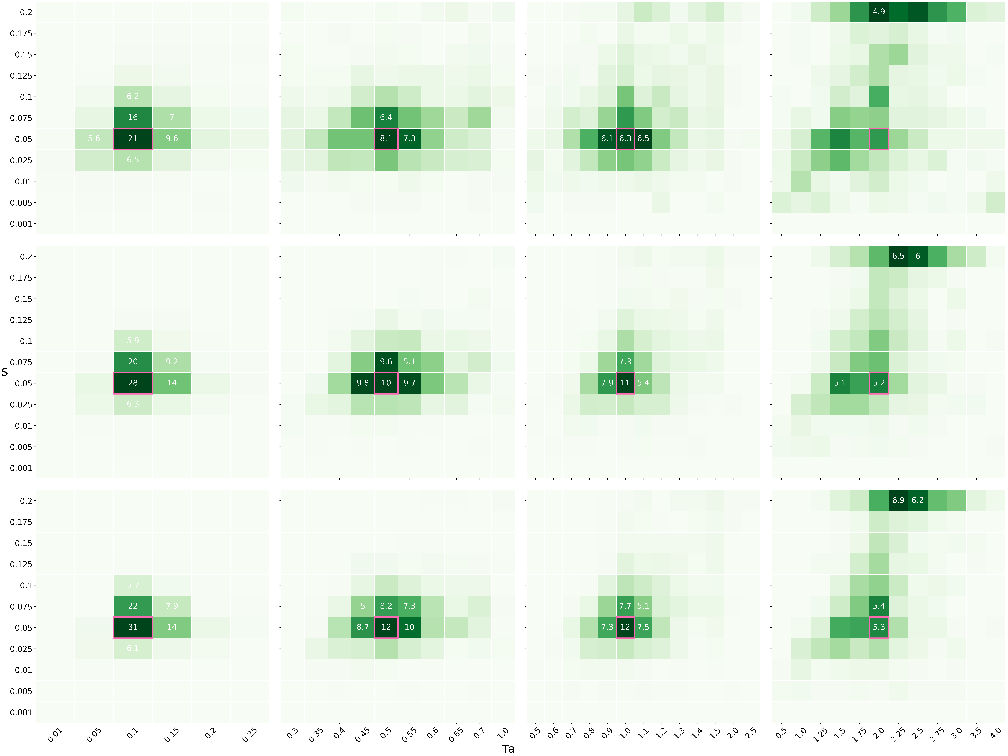
Heatmap, strong selection *s* = 0:05. Results of the gridded optimization. Panels represent different sweep ages (*T_a_* = 0:1; 0:5; 1:0; 2:0 from left to right). Numbers show the percentage of replicates (> 4:5%) associated with a particular parameter combination. The true parameter combination is indicated by a pink square.

**Figure S3:**
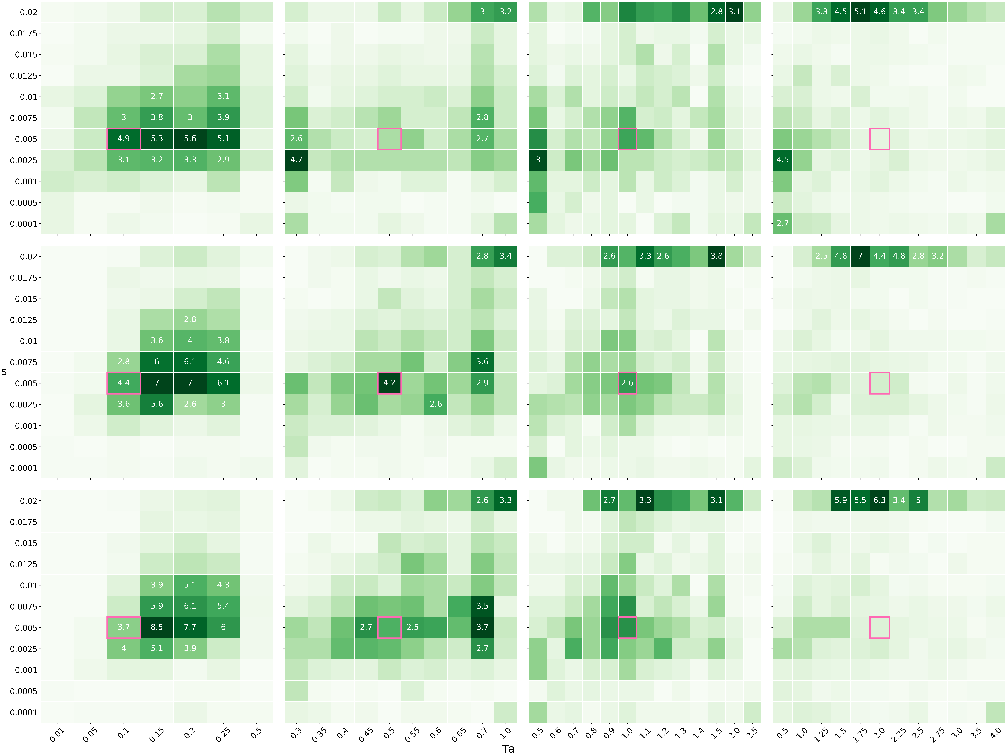
Heatmap, weak selection *s* = 0:005. Results of the gridded optimization. Panels represent different sweep ages (*T_a_* = 0:1; 0:5; 1:0; 2:0 from left to right). Numbers show the percentage of replicates (> 4.5%) associated with a particular parameter combination. The true parameter combination is indicated by a pink square.

**Figure S4:**
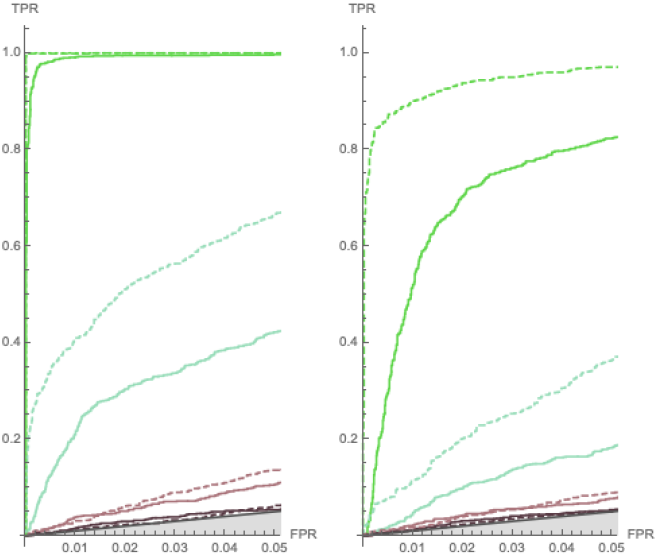
ROC, T_a_=0

**Figure S5:**
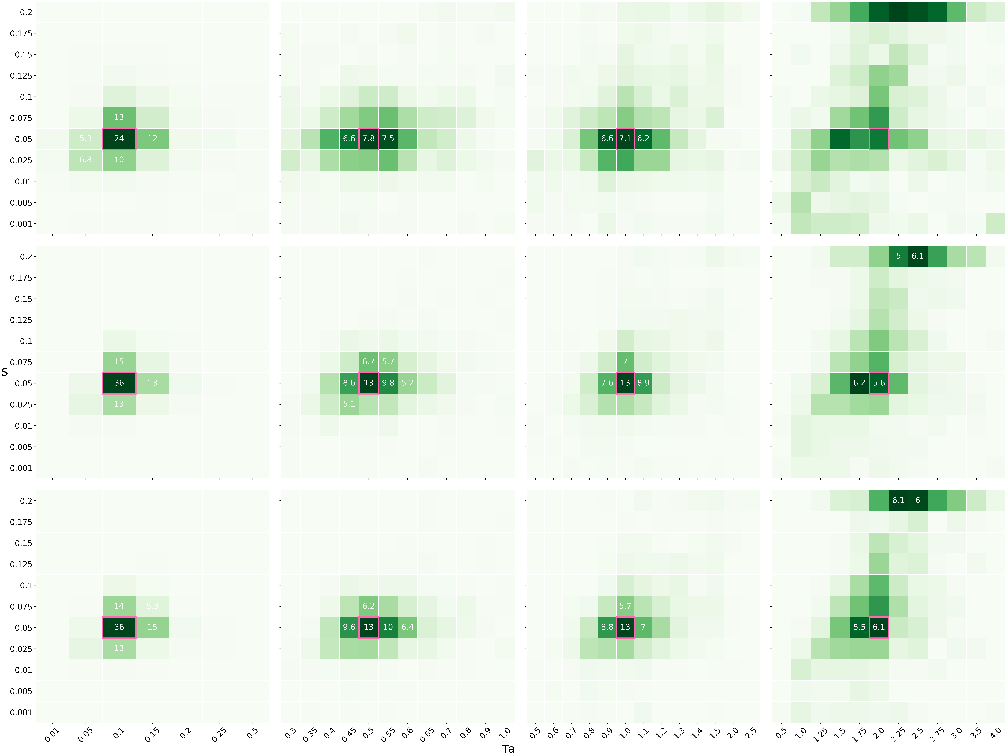
Heatmap, strong selection *s* = 0:05. Results of the gridded optimization. Panels represent different sweep ages (*T_a_* = 0:1; 0:5; 1:0; 2:0 from left to right). Numbers show the percentage of replicates (> 4:5%) associated with a particular parameter combination. The true parameter combination is indicated by a pink square.

**Figure S6:**
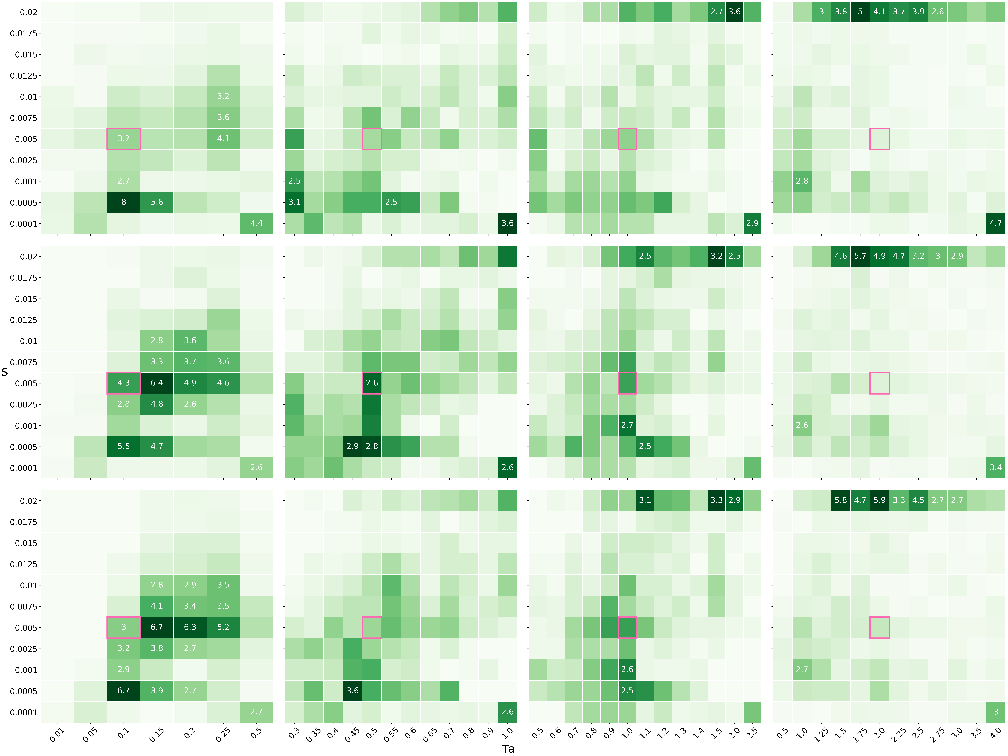
Heatmap, weak selection *s* = 0:005. Results of the gridded optimization. Panels represent different sweep ages (*T_a_* = 0:1; 0:5; 1:0; 2:0 from left to right). Numbers show the percentage of replicates (> 4:5%) associated with a particular parameter combination. The true parameter combination is indicated by a pink square.

**Figure S7:**
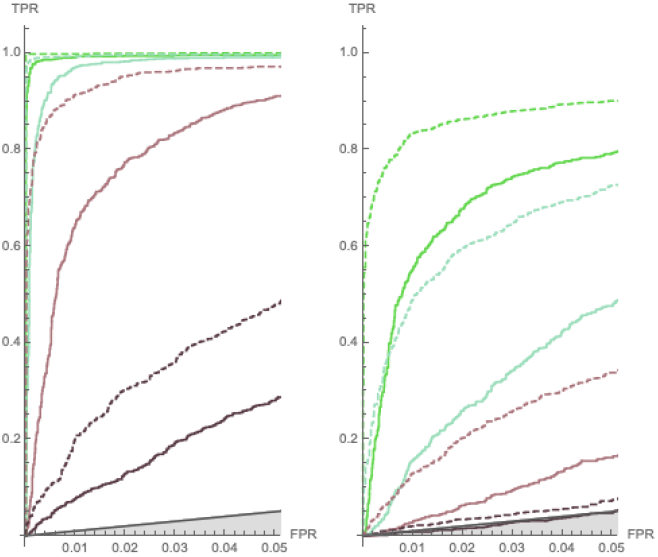
ROC, instantaneous Yule approximation

**Figure S8:**
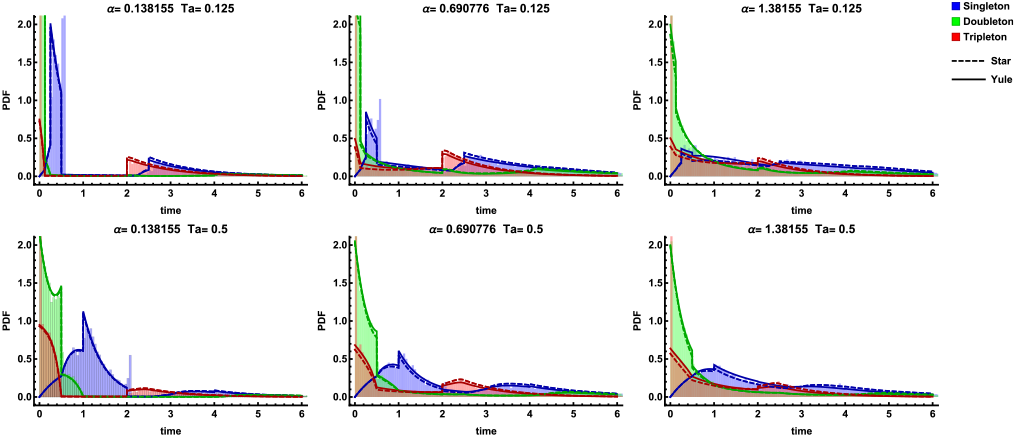
PDF of marginal branch lengths for adaptive introgression scenario. The analytic predictions under the Yule approximation (solid) and star-like approximation (dashed) for a younger sweep (*T_a_* = 0.125, top row) and an older sweep (*T_a_* = 0.5,bottom row) at various distances (*α* ≈ {0.14,0.69,1.4}) from the sweep center. In each panel, the corresponding histogram obtained from 10, 000 replicate simulations is overlaid for comparison. Singleton, doubleton, and tripleton branches are indicated by the colors blue, green, and red, respectively. Here, the time since divergence *T_d_* at the time of the introgression event is 2.0 in units of 2*N* generations, *N* = 10, 000, s = 0.05, *L* = 50, 000, and *r* = 1*e* – 6.

**Figure S9:**
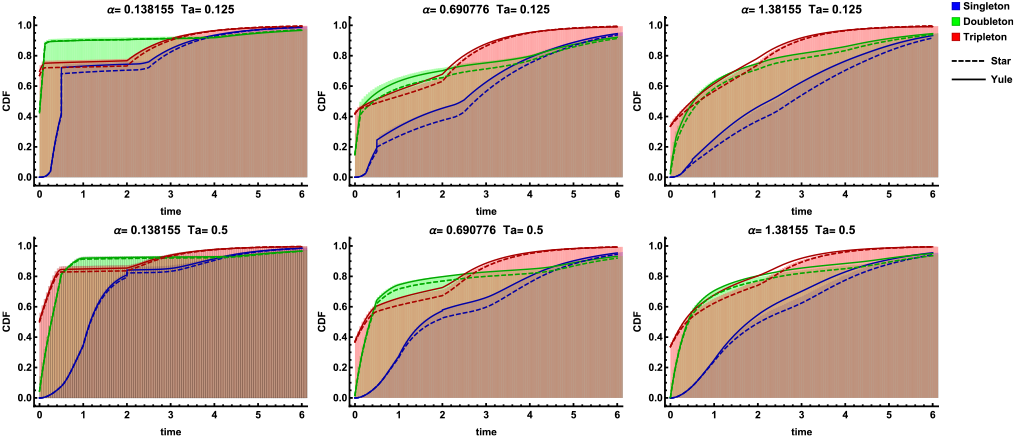
CDF of marginal branch lengths for adaptive introgression scenario. CDF for various distances from the sweep center and times since the sweep. Companion to Fig. S8

